# Influence of Diphenyl Diselenide on Thiol Redox Homeostasis and Electrogenic Membrane Transport in Rotenone-Induced Parkinson’s Disease

**DOI:** 10.1101/2022.10.30.514424

**Authors:** Titilayo Ibironke Ologunagba, Bunmi Olaoluwa Olorundare, Titilope Iyanda, Adenike Adewole, Ige Joseph Kade

## Abstract

The central role of oxidative stress in the etiology of Parkinsons disease defines a key therapeutic role for antioxidant compounds in the management of the disease. Redox-sensitive proteins such as the Na^+^/K^+^-ATPase have also been implicated as one of the targets of oxidative stress. The present study sought to investigate the role of diphenyl diselenide (DPDSe) in amelioration of disturbed redox homeostasis and modulation of enzyme activity caused by rotenone administration to Wistar albino rats. In determining the best route of rotenone administration, animals were grouped into four namely: control, oral, intraperitoneal (IP) and subcutanoeus (SC) and administered rotenone (3mg/kg) via oral, intraperitoneal (IP) and subcutaneous routes, with controls receiving the vehicle (2% DMSO + 98% normal saline (0.85%)). Having observed a more deleterious impact in the with he IP route, this mode of rotenone administration was selected along with oral administration of DPDSe (10mg/kg). this was done with four groups of animals namely: control, DPDSe, rotenone and DPDSe+rotenone. The effect of treatment was evaluated after seven days for total and non-protein thiol levels, lipid peroxidation and Na^+^/K^+^-ATPase activity. The result demonstrated the antioxidant potential of DPDSe in attenuating depletion of thiols, and lipid peroxidation caused by rotenone. It is apparent that DPDSe is a promising therapeutic agent in the management of PD, hence further investigations into its impact on different pathways is expedient in the search for an effective treatment for PD.

**Highlights:**

- Intraperitoneal administration of rotenone gives a comparatively faster development of Parkinson’s disease features in rodents
- Rotenone mediates depletion of total and non-protein thiol levels in the pathogenesis of Parkinson’s disease
- Diphenyl diselenide significantly attenuates thiol depletion and lipid peroxidation mediated by rotenone
- Rotenone-mediated inactivation of Na^+^/K^+^-ATPase in Parkinson’s disease etiology may involve ATP depletion

## INTRODUCTION

There is compelling evidence from literature that oxidative stress may play a central role in the pathogenesis of Parkinson’s disease (PD) (Radak *et al*., 2011; Puspita *et al*., 2017; Poewe *et al*., 2017). Despite the essential physiological roles of reactive oxygen species (ROS) in mediating some signaling pathways as well as cellular response to growth hormones, a failure in the antioxidant machinery in regulating their levels result in oxidative stress. The random oxidation of macromolecules such as proteins, lipids and DNA damage cellular organelles (such as the mitochondria) and structures (such as membrane architecture) and lead to eventual cell death (Wiseman and Halliwell, 1996; Rego and Oliviera, 2003; Sies and Jones, 2017). Parkinson’s disease is typically characterized by the progressive loss of dopaminergic neurons of the substantia nigra pars compacta (SNpc), resulting from a myriad of oxidative stress-related intracellular events, such as disturbed protein homeostasis and mitochondrial dysfunction (Anglade *et al*., 1997; Stefanis, 2005; Alvarez-Erviti *et al*., 2010; Poewe *et al*., 2017).

Evidence from post mortem analysis of the brain of PD patients as well as experimental data linking oxidative stress intricately with mitochondrial dysfunction is convincing (Parker *et al*., 2008; Poewe *et al*., 2017; Puspita *et al*., 2017). The mitochondria are the major site of ROS production, target of oxidative assault especially by PD-inducing neurotoxins such as rotenone (Cannon and Greenamyre, 2013; Terron *et al*., 2018), 6-hydroxydopamine (Ungerstedt *et al*., 1974; Hwang, 2013). Rotenone is a naturally-derived pesticide extracted from plant roots used as household pesticide and used for eradicating nuisance fish populations in lakes and reservoirs (Betarbet *et al*., 2000). This compound is highly hydrophobic and can readily cross the blood brain barriers and biological membranes independent of the dopamine transporter (DAT). It is also a classical, well-characterized, specific inhibitor of mitochondrial complex-I (Thiffault *et al*., 2000; Radad *et al*., 2006). The inhibition of mitochondrial complex-I culminates in mitochondrial dysfunction, ROS production and ATP depletion (Cannon and Greenamyre, 2013; Terron *et al*., 2018).

The depletion of ATP significantly alters the function of ATP-dependent cellular processes, such as the maintenance of electrochemical gradient of Na^+^ and K^+^ ions across the plasma membrane, which is coordinated by the Na^+^/K^+^-ATPase (Skou, 1957, Kaplan, 2002). The Na^+^/K^+^-ATPase is sulfhydryl, transmembrane integral protein and is vulnerable to inhibition by disturbed redox homeostasis (Omotayo *et al*., 2015; Bogdanova *et al*., 2016), hence a primary target of oxidative stress. In fact, the involvement of inhibition of this protein as a component of the pathogenesis of some degenerative diseases such as Alzheimers disease and PD is being widely explored (Shrivastava *et al*., 2015; Dichiara *et al*., 2017; Diggelen *et al*., 2019; Bejcek *et al*., 2021). Hence, the search for therapeutic interventions for PD management have widely explored the promising potential of antioxidant compounds.

Specifically, several reports have demonstrated the ameliorative effects of antioxidant compounds in ameliorating the pathological features of PD (Kujawska and Jodynis-Liebert, 2018; Costa *et al*., 2016; Moosavi *et al*., 2016; Pan *et al*., 2003). Polyphenols such as quercetin (Karuppagounder *et al*., 2013; Haleagrahara *et al*., 2011), kaempferol (Li and Pu, 2011), rutin (Khan *et al*., 2012), hesperidin (Antunes *et al*. 2014), among others, have demonstrated significant degrees of potency in ameliorating the pathology of PD. One of the neuroprotective effects associated with these compounds in their ability to improve the thiol redox status of the test animals by elevation of glutathione (GSH) levels (Lv *et al*. 2012: Antunes *et al*. 2014), elevation of glutathione peroxidase (GPx) activity (Antunes *et al*., 2014; Lv *et al*., 2012; Li and Pu, 2011), reduction of thiobarbituric acid reactive substances (TBARS) (Khan *et al*., 2012) as well as elevation of Na^+^/K^+^-ATPase activity (Lv *et al*., 2012).

The upregulation of thiol redox status has been attributed to a class of synthetic antioxidant compounds, the organochalcogens. These compounds exhibit excellent GPx mimetic properties and have demonstrated potent antioxidant properties in different oxidative stress-linked disease models (Zamberlan *et al*., 2014; Kade *et al*., 2008, 2009; Nogueira and Rocha, 2010; Kade, 2016). Diphenyl diselenide (DPDSe), an organochalcogen, has gained wide attention for its antioxidant properties. In different animal models of oxidative stress, the thiol replenishing ability of DPDSe has been demonstrated, along with its ability to protect the sulfhydryl, redox sensitive Na^+^/K^+^-ATPase from oxidative assault (Kade *et al*., 2008; Kade, 2016). It is also being explored for its therapeutic potential in Parkinson’s disease models (Sampaio *et al*., 2017; daRocha *et al*., 2013). However, it is not clear whether the protective effect of DPDSe on Na^+^/K^+^-ATPase is involved in its therapeutic potential in Parkinson’s disease, hence the present study.

## MATERIALS AND METHOD

### Materials

rotenone, 5’5’-dithio-bis (2-nitrobenzoic) acid (DTNB), diphenyl diselenide (DPDSe), adenosine triphosphate (ATP), ouabain, glycine, dimethyl sulphoxide (DMSO) were obtained from Sigma-Aldrich (St. Louis, MO USA). All other chemicals of analytical grade were obtained from standard commercial suppliers

### Animals

Male adult Wistar rats (200–250g) obtained from our own breeding colony at Biochemistry Department (FUTA) animal house were used. Animals were kept in separate animal cages, on a 12-h light: 12-h dark cycle, at a room temperature of 22–24°C, and with free access to food and water. The animals were used according to standard guidelines on the Care and Use of Experimental Animal Resources (National Research Council, 2011).

### Parkinson’s disease induction

The best route of rotenone administration was first determined. Herein, a 50X stock solution of rotenone dissolved in 100% DMSO was prepared and stored for the preparation of daily fresh dilutions to a final concentration of 3mg/kg rat weight in a solution of 2% DMSO + 98% normal saline (0.85%) via oral, intraperitoneal and subcutaneous routes of administration. In this randomized experiment, sixteen male Wistar albino rats (150-200) were grouped into four groups of four animals each and treated for seven days: (group 1) control; (group 2) rotenone (3 mg/kg) oral route; (group 3) rotenone (3 mg/kg) intraperitoneal route; (group 4) rotenone (3 mg/kg) subcutaneous route. Animals were fasted twelve (12) hours prior to euthanasia.

### Treatment

From the first experiment, intraperitoneal route of administration demonstrated the most impact, hence rotenone administration via the IP route was selected. Sixteen male Wistar albino rats (150-200) were randomly grouped into the following groups: (group 1) control; (group 2) DPDSe (10 mg/kg) alone; (group 3) rotenone (3 mg/kg) intraperitoneal route; (group 4) DPDSe (10 mg/kg) + rotenone (3 mg/kg) intraperitoneal route. Treatment by oral administration of DPDSe was administered thirty (30) minutes prior to rotenone treatment, for seven days. After expiration of the duration of treatment, animals were fasted 12 hours prior to euthanasia. Thereafter, they were euthanised and sacrificed by decapitation. The brain tissue was quickly removed, dissected into five brain regions, homogenized and used for biochemical assays of redox parameters (TBARS and total and non-protein thiol levels) and enzyme activity.

### Tissue preparation

Animals were decapitated and the whole brain tissues removed. The brain tissue was sectioned into the regions of interest, cortex, cerebellum, midbrain, striatum and hippocampus. The sections were individually weighed and homogenized in cold 50 mM Tris–HCl pH 7.4. The homogenate was centrifuged at 4,000 g for 10 min to yield the low-speed supernatant (S1) fraction that was used for biochemical assays. For all analyses, protein content was determined by the method of Lowry *et al*. (1951), using bovine serum albumin as the standard.

### Assay of Total and Non-protein thiols

Total and non-protein thiol levels were evaluated after deproteinization with TCA (5% in 1 mmol/EDTA) according to the method described by Ellman (1959).

### Assay of TBARS

Production of TBARS was determined as described by Ohkawa *et al*. (1979) except that the buffer for the color reaction was pH 3.4. The color reaction was developed by adding 300 μl 8.1% SDS to S1, followed by sequential addition of 500 μl acetic acid/HCl (pH 3.4) and 500 μl 0.8% TBA. This mixture was incubated at 95°C for 1 h. TBARS produced were measured at 532 nm, and the absorbance was compared to that of a standard curve obtained using malondialdhyde (MDA).

### Assay of Na^+^/K^+^-ATPase

Cerebral Na^+^/K^+^-ATPase activity was measured as described by our group (Kade *et al*., 2008) except that there was no 10 min pre-incubation. Phosphate was measured according to Fiske and Subbarow (1925).

### Behavioural studies

#### Open field test

Rats were individually placed in a wooden cage square measuring 60 cm×60 cm divided into sixteen squares using a marker. The numbers of crossing made across the internal squares as well as rearing (upright posture) were recorded over a period of 5 minutes.

#### Hanging wire test

Rats were individually placed on 2mm thick metallic wire that is secured to the vertical stands of a wooden frame. The rats catch the wire with their limbs and remain suspended on the wire for a duration of 5 minutes. The number of falls of each animal within this time frame is recorded.

### Statistical analysis

All experiments were performed in triplicate and data collected were analysed statistically by ANOVA. These were followed by Duncan’s new multiple range test where appropriate. All differences with p<0.05 will be considered significant.

## RESULTS-pg 94, thesis

### Determining the effect of different routes of rotenone administration

#### Behavioral studies

As presented in Figure 1 (panel A), all three groups of rotenone -treated animals displayed a significant loss of body weight compared to the control group. However, there was a varying degree of weight loss for individual routes of rotenone administration. Test animals in the oral rotenone administration group showed the least weight loss while the animals in the intraperitoneal (IP) rotenone administration groups showed the most significant weight loss after the 7-day period of treatment (p < 0.05). This was accompanied with 25% mortality in test animals. Furthermore, the development of motor symptoms following rotenone adminsitration as evaluated by hanging wire test (Figure 1, panel B) and open-field test (Figure 1, panels C and D) of animals show a similar pattern with the observation of weight loss. Herein, one-way ANOVA (different routes of administration) of the result presented in Figure 1, panel B revealed that test animals in the IP group had more falls (p < 0.05) compared to the control group as well as the two other route of administration [oral and subcutaneous (SC)]. In addition, open-field test was used to investigate the rearing behavior of test animals following treatment. As presented in Figure 1, panel C in the intraperitoneal (IP) route group showed less rearing (panel C) as they appeared very weak and less interested. More so, the average number of crossings of internal squares (panel D) within the cage was significantly less for IP rotenone group. Taken together, the behavioural studies highlight that intraperitoneal route of administration may be the best route for rotenone administration for the subsequent study.

**Figure 1:**
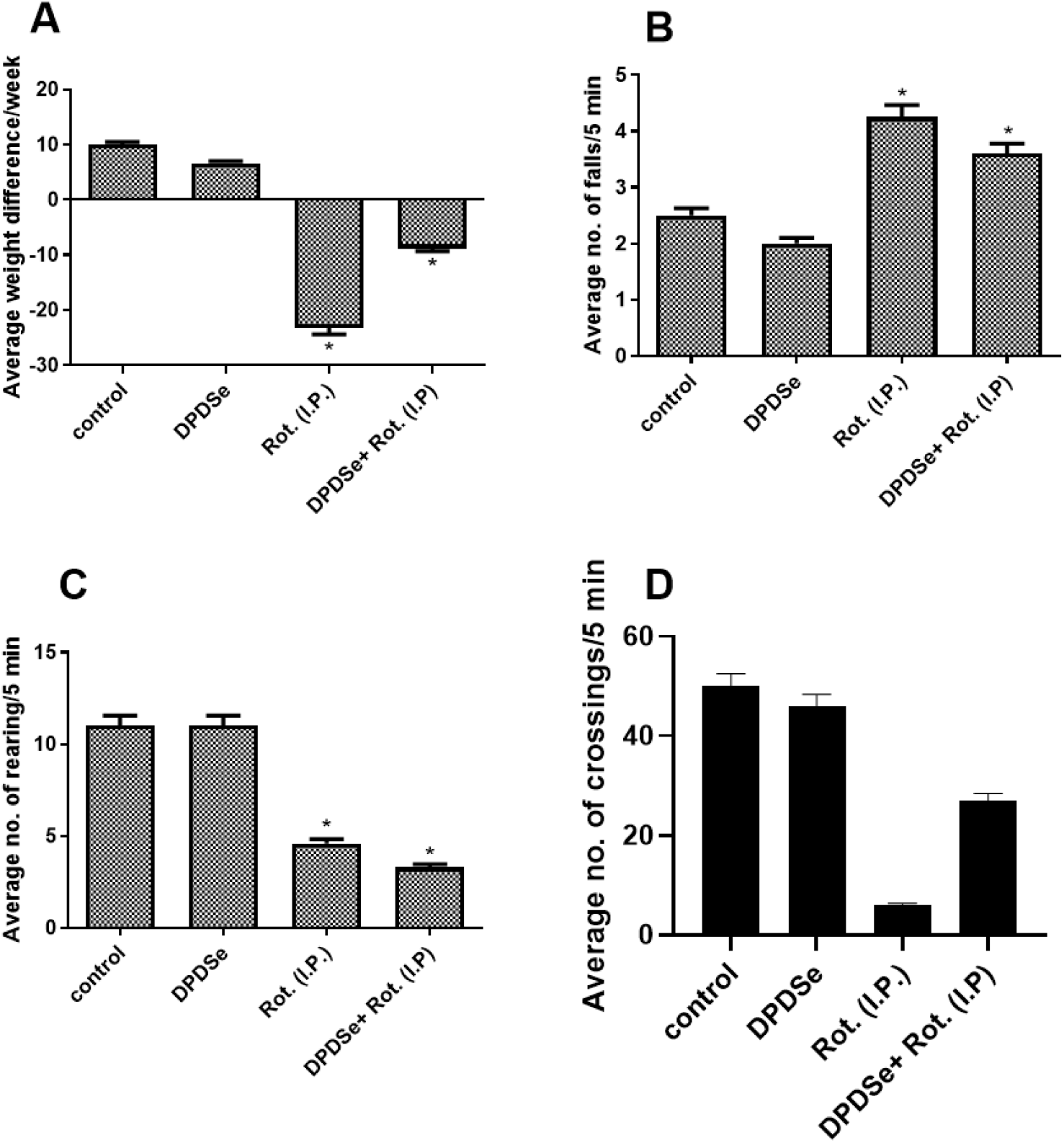
Average weight difference (panel A) in seven days, hanging wire test (panel B), open field test-rearing (panel C) and crossing (panel D) within five minutes of observation in a 60cm x 60cm wooden square cage, for indicated routes of rotenone (3mg/kg) administration. Data were presented as mean ± SEM of at least three experimental animals carried out on different days. * indicates significantly lower than control (p < 0.05).

### Effect of rotenone on cellular redox status-total and non-protein thiol levels

The impact of rotenone on redox status of test animals is here assessed by a measure of total and non-protein thiol levels, in five rain regions, the cortex, cerebellum, midbrain, striatum and hippocampus. One-way ANOVA of the result of total protein thiol levels (Figure 2) showed rotenone significantly depleted the levels of total and non-protein thiols in the cortex, and this is more is more prominent in the I.P group (p < 0.05). In the cerebellum, one-way ANOVA of the result presented in panel D showed a more significant depletion of non-protein thiols than was observed in the cortex (panel B), while the results for the midbrain (panels E and F) showed a more significant depletion of total protein thiols than was observed in the cortex and cerebellum. Furthermore, the striatum (panels G and H) and the hippocampus (panels I and J) showed a similar trend of non-protein thiol depletion in the rotenone treated groups as seen in the hippocampus, where the oral route of administration appeared to cause less total and non-protein thiol depletion. Summarily, the I.P route of administration caused more significant depletion of total and non-protein thiols in all five regions of the brain investigated indicating disturbed thiol redox homeostasis.

**Figure 2:**
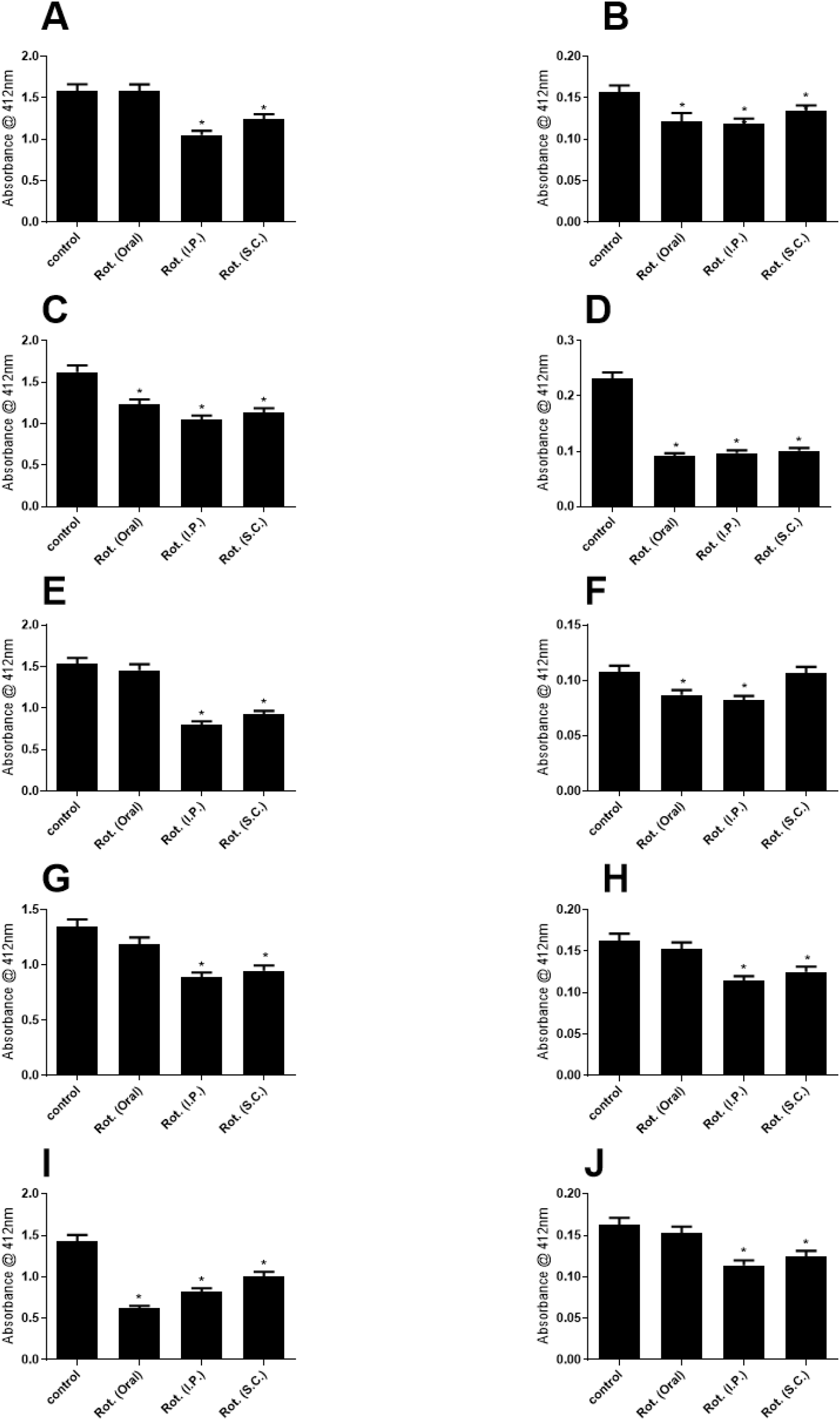
Estimation of total thiol and non-protein thiol in the cortex (panels A and B), cerebellum (panels C and D), midbrain (panels E and F), striatum (panels G and H) and hippocampus (panels I and J) respectively *in vivo* for oral, intraperitoneal and subcutaneous routes of rotenone (3mg/kg) administration. Data are presented as mean ± SEM for independent experiments done in duplicate carried out in different days. * represent significant difference from control at p < 0.05.

### Effect of rotenone on lipid peroxidation and activity of Na^+^/K^+^-ATPase

Since the previous section demonstrated the disturbance of thiol redox status by rotenone, it is necessary to evaluate the extent of lipid peroxidation and modulation of the activity of the redox-sensitive Na^+^/K^+^-ATPase. In the cortex (panels A and B), one-way ANOVA (p < 0.05) of the results showed that lipid peroxidation was significantly elevated by rotenone via all three routes of administration. This was accompanied by simultaneous inhibition of Na^+^/K^+^-ATPase which was more significant in the I.P route. This result is in correlation with the depletion of total and non-protein thiols presented in the previous section. Furthermore, the cerebellum (panels C and D) and hippocampus (panels I and J) appear to have a similar trend of the effect of rotenone administration on TBARS and Na^+^/K^+^-ATPase activity and in agreement with the events described in the cortex. In the midbrain (panels E and F), it was observed that lipid peroxidation and Na^+^/K^+^-ATPase inhibition was almost at the same level for all three routes, while the striatum (panels G and H) presents a likelihood of more lipid peroxidation (panel A) by subcutaneous route of rotenone administration when compared to the I.P route. This is a slight deviation from what has been observed so far. However, it is clear that rotenone administration via all three routes significantly induced peroxidation of lipids and altered activity of the Na^+^/K^+^-ATPase.

### Investigating the neuroprotective effects of DPDSe against rotenone assault

The antioxidative effects of DPDSe is widely reported, hence in the present study, its ameliorative effect on rotenone (3mg/kg body weight i.p)-mediated modulation of redox homeostasis and enzyme activity was investigated.

#### Behavioural studies

The assessment of body weight of test animals revealed a significant (p<0.05) amelioration of rotenone-induced weight loss by DPDSe treatment (Figure 4, panel A). The fine motor skills of test animals tested by the hanging wire test is presented in panel B. Here, DPDSe apparently improved the test animals’ motor skills partially as the number of falls observed with DPDSe-treated animals was comparatively less than observed in the rotenone(i.p)-treated group. Furthermore, the result of open-field test presented in panel C apparently showed that DPDSe had no significant effect on the impairment of rearing behaviour (panel A) or reduced number of crossings (panel B) mediated by rotenone administration (p < 0.05). Taken together, DPDSe had partial effect in the reversal of the deleterious impact of rotenone administration on behavioural parameters investigated in the present study.

### Effect of DPDSe on rotenone-mediated depletion of total and non-protein thiols

The antioxidant potential of DPDSe widely highlighted in literature makes it a worthy candidate in the amelioration of rotenone mediated depletion of total and non-protein thiols in this model. Assessing the five different regions of the brain earlier examined-the cortex, cerebellum, midbrain, striatum and hippocampus-diphenyl diselenide presents some talking points. In the cortex, one-way ANOVA of the result presented in Figure 5 revealed that treatment with DPDSe significantly elevated total (panel A) and non-protein thiol (panel B) levels, especially in rotenone-administered animals. However, in the cerebellum (panels C and D), DPDSe had less effect on restoration of rotenone-mediated depletion of total and non-protein thiols. The midbrain (panels E and F), striatum (panels G and H), and hippocampus (panels I and J) showed significant ameliorative effect of DPDSe on rotenone-mediated disturbed thiol redox homeostasis. It is however, worthy of note that in the midbrain (panels E and F), which is the epicenter of PD pathology, DPDSe significantly elevated total and non-protein thiol levels in the absence of rotenone as compared to the other brain regions investigated. While this is expected, amelioration of total protein thiol depletion by rotenone treatment was apparently minimal (panel E). Further investigation could reveal the import of this observation. Summarily, DPDSe partially ameliorated the effect of rotenone administration on disturbed thiol redox homeostasis.

### Effect of DPDSe on rotenone-mediated lipid peroxidation and enzyme inhibition

The modulation of thiol redox status by DPDSe and rotenone is expected to translate to modulation of lipid peroxidation and activity of the Na^+^/K^+^-ATPase, being a redox-sensitive protein. As presented in Figure 6, the cortex showed reduction of TBARS production (panel A) in DPDSe-treated animals, however, there was no significant recovery of Na^+^/K^+^-ATPase activity (panel B) observed (p < 0.05). in the cerebellum (panels C and D), midbrain (panels E and F) and striatum (panels G and H), rotenone administration significantly elevated TBARS production as compared to what was observed in the cortex (panel A) and hippocampus (panels I and J). DPDSe treatment modulated TBARS levels significantly in the former (cerebellum, midbrain and striatum) however, its effect on the activity of Na^+^/K^+^-ATPase in all five regions showed much less efficiency in the recovery of enzyme activity in rotenone-treated animals.

## DISCUSSION

Parkinson’s disease (PD) is a progressive neurodegenerative disorder characterized by a progressive loss of the dopaminergic neurons of the substantia nigra. Several efforts at unraveling the pathogenesis of this disease have highlighted a dysregulation of redox homeostasis and protein homeostasis (Poewe *et al*., 2017; Hwang, 2013; (Radak *et al*., 2011; Puspita *et al*., 2017). In particular, the modulation of key redox-sensitive proteins have also been reported. One of such is the Na^+^/K^+^-ATPase-a transmembrane enzyme responsible for the maintenance of electrochemical gradient across the plasma membrane (Yu, 2003; de Carvalho *et al*., 2004; Rodacker *et al*., 2006; Aperia, 2007; Shrivastava *et al*., 2015). While the debate about the cause or effect relationship between this enzyme’s inhibition and Parkinson’s disease is ongoing (Shrivastava *et al*., 2015; Dichiara *et al*., 2017), it is undisputed that this crucial enzyme plays an important role in the etiology of Parkinson’s disease, given its physiological relevance. The Na^+^/K^+^-ATPase is also implicated in a number of oxidative stress-related degenerative diseases, for which therapeutic intervention by antioxidant compounds is being explored (Kade *et al*., 2008, 2009). One of such promising antioxidant compounds is diphenyl diselenide (DPDSe). DPDSe is known to display GPx-mimetic properties in its antioxidant action hence its suitability to amelioration of oxidative stress. While the therapeutic potential of DPDSe in Parkinson’s disease is being investigated (daRocha *et al*., 2013; Sampaio *et al*., 2017), its effect on the activity of Na^+^/K^+^-ATPase in this condition has not been reported. Hence, the present study sought to investigate this relationship using rotenone model of Parkinson’s disease *in vivo*.

Rotenone is a highly lipophilic compound that has displayed excellent capacity to cross the blood brain barrier. The observed variation in presentation of PD-like symptoms with different routes of administration, in the present study, further highlights previous observations about rotenone metabolism. Rotenone is a specific inhibitor of mitochondrial complex I. However, it is poorly absorbed from the gastrointestinal (GI) tract (Fang and Casida, 1998; Caboni *et al*., 2004). Hence, the observed minimal effect of rotenone in the oral administration group is likely due to its poor absorption via this route (Figure 1, panel A). It has also been reported that approximately 20% of orally administered rotenone is excreted via the urine 24h after treatment, and unabsorbed rotenone in the GI tract is excreted through the faeces (Caboni *et al*., 2008; Gupta 2012). On the other hand, the IP route showed the most significant effect of rotenone on weight loss, motor coordination as well as rearing and crossing behaviour of test animals. This is likely related to the presence of extensive perfusion of the peritoneum with blood capillaries, favouring an excellent surface area for exchange of xenobiotics between the peritoneal cavity and blood plasma (Aune *et al*., 1970; Turner *et al*., 2011; Al Shoyaib *et al*., 2020). This observation was further verified by the biochemical assays of the redox status and activity of the Na^+^/K^+^-ATPase in brain regions associated with motor functions.

The observed significant depletion of total and non-protein thiols in all five regions of the brain investigated suggests that the development of PD features is accompanied (or preceded) by a disturbance of the thiol redox status in the brain (Figure 2). However, since animals intraperitoneally injected with rotenone consistently showed the most significant total and non-protein thiol depletion, it is clear that this route is best suited for the time-course of treatment (7 days). Hence, a significant disturbance of thiol homeostasis is a precursor to oxidative stress induction in the dopaminergic neurons of the substantia nigra. The observance of a measure of total and non-protein thiols depletion for all three routes of rotenone administration (oral, I.P and subcutaneous), suggests that disturbance of thiol homeostasis precedes the development of the PD features observed in the behavioural studies. This is likely a component of the pathogenesis of Parkinson’s disease.

Disturbance of thiol redox status may be correlated with lipid peroxidation and inactivation of redox-sensitive proteins such as the Na^+^/K^+^-ATPase since depletion of the cell’s primary antioxidant machinery (thiols) leads to increased ROS levels. Membrane lipids and redox-sensitive proteins are primary targets of oxidative stress as earlier discussed. In the present study, observed significant elevation of lipid peroxidation and enzyme inhibition in rotenone-treated animal (p<0.05) further buttresses previous reports (Figure 3) (Braak *et al*., 2003; Poewe *et al*., 2017). It is apparent from the present study that there is a prominent role for the enteric nervous system (ENS) in the propagation of disease features as widely reported. This is because the IP route of administration displayed a significant potential in disease induction. In fact, in explaining the basis of ENS neurodegeneration in rotenone-treated mice, it has been described as the portal for α-synuclein transmission to the central nervous system (CNS), as well as basis for gastrointestinal (GI) symptoms in early PD (Drolet *et al*., 2009; Greene *et al*., 2009; Pan-Montojo *et al*., 2012, 2012). Furthermore, enteric neurons as being similarly vulnerable as nigrostriatal neurons, having unmyelinated axons and multiple synaptic endings. Thus increasing their vulnerability in PD and facilitating the uptake and transmission of α-synuclein through their connecting ganglia (Paillusson *et al*., 2013; Sharrad *et al*., 2017; Al Shoyaib *et al*., 2020).

**Figure 3:**
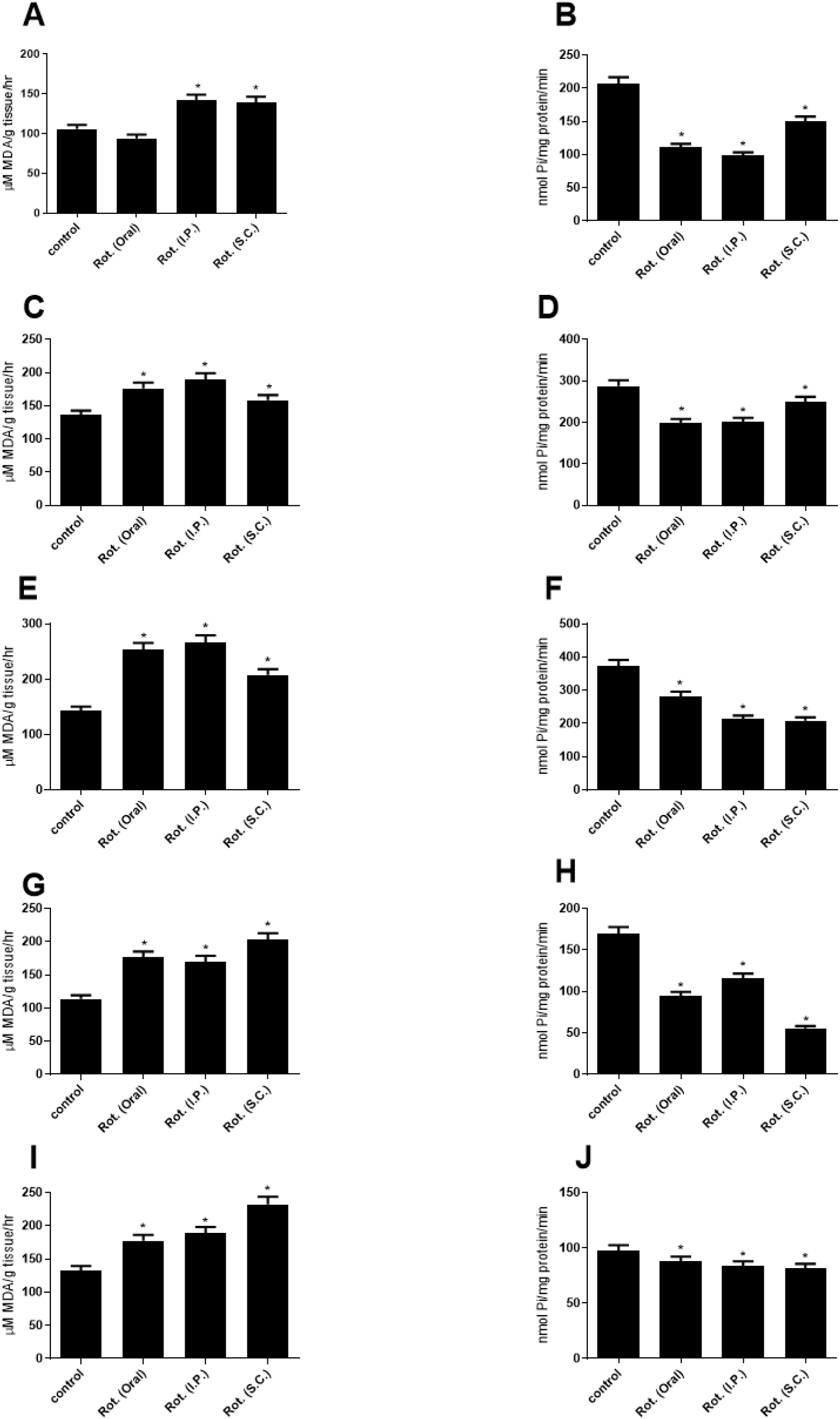
Estimation of lipid peroxidation and Na^+^/K^+^-ATPase activity respectively in cortex (panels A and B), cerebellum (panels C and D), midbrain (panels E and F), striatum (panels G and H) and hippocampus (panels I and J) *in vivo* for oral, intraperitoneal and subcutaneous routes of rotenone (3mg/kg) administration. Data are presented as mean ± SEM for independent experiments done in duplicate carried out in different days. * represent significant difference from control at p < 0.05.

**Figure 4:**
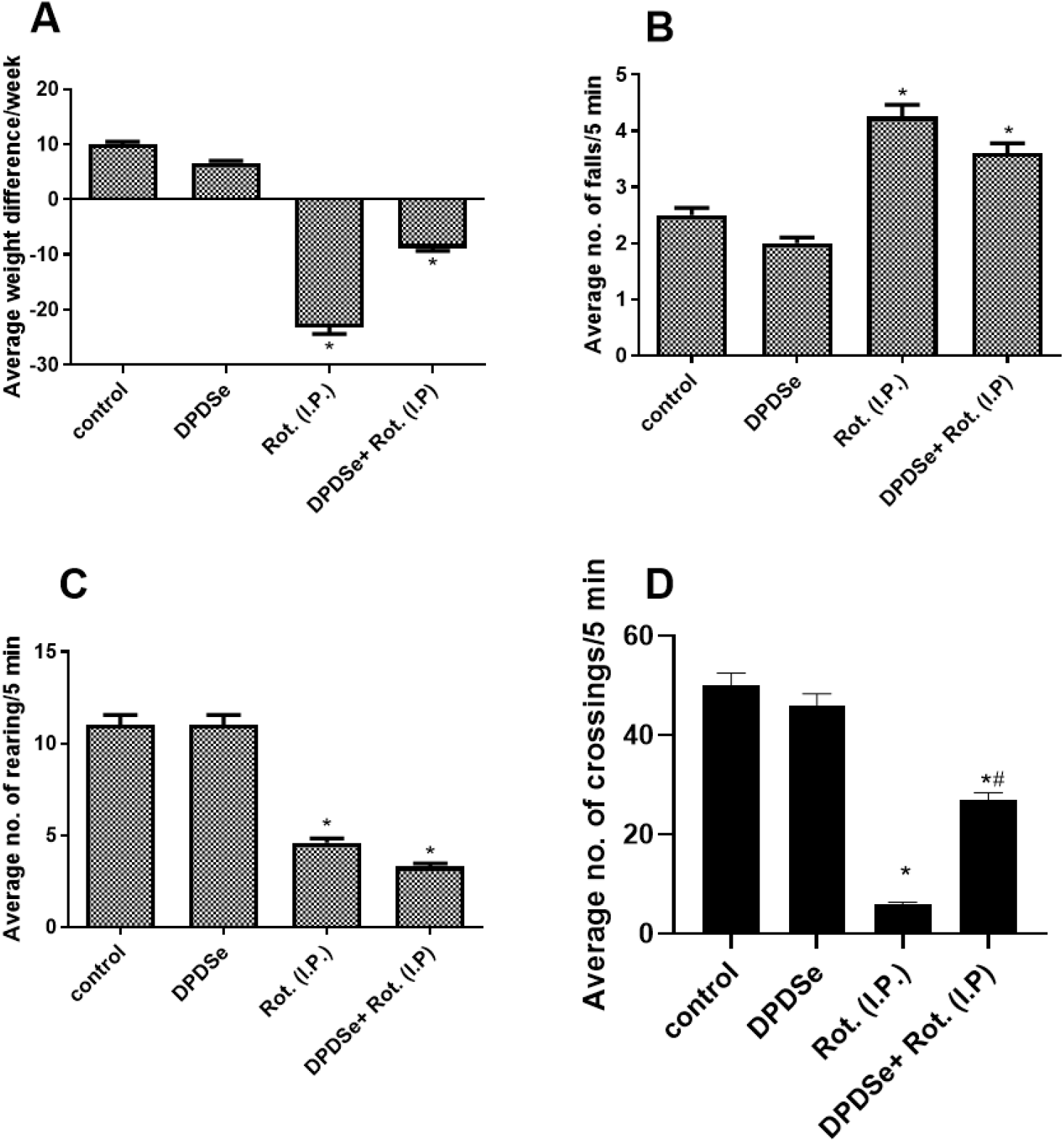
Average weight difference (panel A) in seven days, hanging wire test (panel B), open field test-rearing (panel C) and crossing (panel D) within five (5) minutes of observation in a 60cm x 60cm wooden square cage, for indicated treatment conditions. Data were presented as mean ± SEM of at least three experimental animals carried out on different days. * indicates significantly lower than control (p < 0.05).

**Figure 5:**
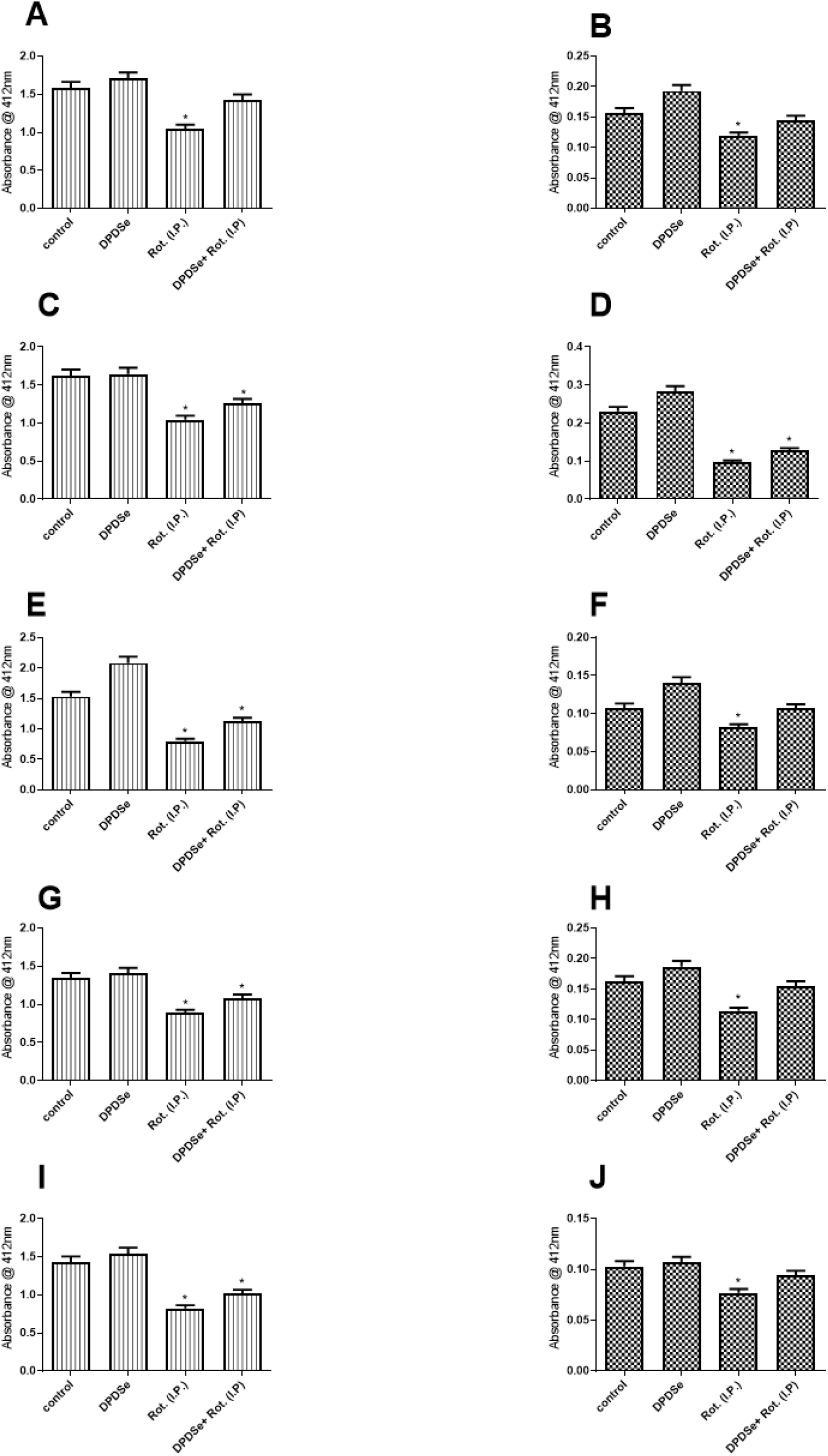
Estimation of total thiol and non-protein thiol in cortex (panels A and B), cerebellum (panels C and D), midbrain (panels E and F), striatum (panels G and H) and hippocampus (panels I and J) *in vivo* for indicated treatment conditions. Data are presented as mean ± SEM for independent experiments done in duplicate carried out in different days. * represent significant difference from control at p < 0.05.

**Figure 6:**
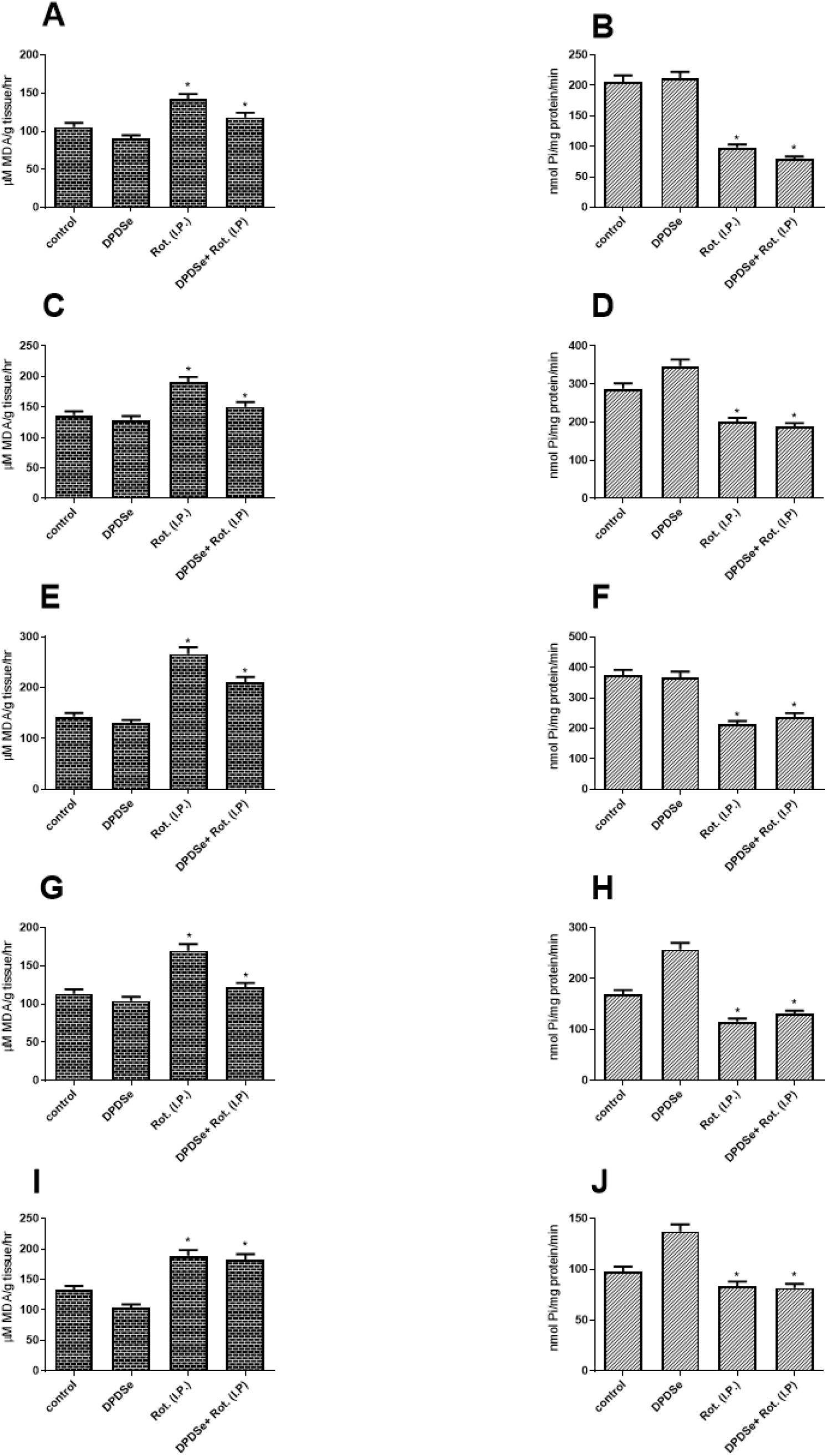
Estimation of lipid peroxidation and Na^+^/K^+^-ATPase activity in cortex (panels A and B), cerebellum (panels C and D), midbrain (panels E and F), striatum (panels G and H) and hippocampus (panels I and J) *in vivo* for indicated treatment conditions. Data are presented as mean ± SEM for independent experiments done in duplicate carried out in different days. * represent significant difference from control at p < 0.05.

More so, the absorption of substances via the intraperitoneal route have been associated with the physicochemical properties of the substance, wherein a lipophilic substance is more rapidly absorbed into tissues and cleared from systemic circulation following IP administration (Canal *et al*., 1989). In comparison to SC route, different reports have demonstrated a higher absorption rate and systemic circulation of drugs via the IP route that the subcutaneous (SC) route (Lee *et al*., 2011; Zhang *et al*., 2014; Wrangler *et al*., 2016; Sumbria *et al*., 2013; Statler *et al*., 2007; Parkes *et al*., 2001, Fam *et al*., 2013; Shi *et al*., 2012; Gerrard *et al*., 1992). It is thus clear that the IP route of rotenone administration is an excellent model in the study of PD as well as therapeutic intervention by antioxidants.

The excellent antioxidant properties of diphenyl diselenide previously reported in literature was here explored (Kade *et al*., 2008, 2009; Nogueira and Rocha, 2010; Kade, 2016). Its ability to improve thiol redox levels, ameliorate lipid peroxidation as well as relieve the inhibition of sulfhydryl proteins such as the Na^+^/K^+^-ATPase in vivo (Kade *et al*., 2008, 2009) was further buttressed in the present study. The ameliorative effect of DPDSe on weight loss and motor symptoms (Figure 4) mediated by rotenone further asserts the therapeutic potential of this compound.

In agreement with previous reports (Kade *et al*., 2008; Kade, 2016), DPDSe significantly replenished total and non protein thiols depleted by rotenone (Figure 5) Thus demonstrating the potential of DPDSe to protect dopaminergic neurons from oxidative assault. Taken together, DPDSe improved the thiol redox status of all five regions of the brain. This forms a critical basis for the observed ameliorative effect of DPDSe on lipid peroxidation in the brain regions investigated (Figure 6). Disturbed thiol redox status poses a major threat to redox balance in the neurons as well as the function of redox-sensitive proteins such as the Na^+^/K^+^-ATPase. This redox-sensitive electrogenic pump is critical to signal transduction at the synaptic clefts (Shrivastava *et al*., 2018) and its inhibition could lead to disruption of signal transduction and depolarization of nerve endings (Lees, 1991; Aperia, 2007). Moreover, rotenone is a specific inhibitor of mitochondrial complex I. Thus, inhibition of mitochondrial complex I by rotenone could lead to impaired ATP production and increased ROS production (Aperia *et al*., 2016; Shrivastava *et al*., 2017; 2018). Hence, enzyme inhibition is not only correlated with increased TBARS levels and altered thiol redox status, but also to the possibility that ATP depletion (as a result of rotenone-mediated inhibition of mitochondrial complex-I) further contributes to enzyme inhibition. It is thus obvious that a myriad of intracellular events are associated with rotenone toxicity, and rotenone-mediated mitochondrial dysfunction is critically connected with reduced Na^+^/K^+^-ATPase activity.

It is apparent that a plethora of molecular events culminate into the pathogenesis and progression of PD. Further investigation into the effect of DPDSe in other PD-associated molecular pathways or proteins is expedient in the appreciation of its therapeutic potential. Another study on the therapeutic potential of DPDSe highlighted its attenuation of increased mechanical and thermal nociception caused by 6-hydroxydopamine administration (daRocha *et al*., 2013). Future perspectives in appreciation of the therapeutic potential could take into of its lipophilicity as well as the contribution of the enteric nervous system in the pathogenesis of PD to optimize this promising antioxidant compound.

## Funding

This research was partly funded by the DBT-TWAS Postgraduate Fellowship (GCU-4/DBT-TWAS/105/#18777) awarded to TIO.

## Acknowledgement

TIO and IJK acknowledge the financial support of DBT-India and TWAS. The authors also acknowledged the financial support of CAPES, FINEP, FAPERGS, PRONEX and CNPq, FINEP research grant Rede Instituto Brasileiro de Neurociencia (IBN-Net) # 01.06.0842-00 and the INCT for excitotoxicity and Neuroprotection-CNPq and CNPq-ProAfrica grant awarded to IJK and JBTR.

## RESULTS

## Notes

### Competing Interest Statement

The authors have declared no competing interest.

